# A standardized toolkit for programmable protein secretion and surface display in *Bacteroides thetaiotaomicron*

**DOI:** 10.64898/2026.05.12.724727

**Authors:** Yu-Hsuan Yeh, Shannon J. Sirk

## Abstract

*Bacteroides thetaiotaomicron* (*Bt*), a dominant bacterial species in the human gut, is a promising chassis for engineering in situ therapeutic delivery systems. Previously, we developed a secretion toolkit composed of endogenous lipoprotein signal peptides (SPs) and full-length secretory proteins from *Bt*. However, due to variations in length, structure, and amino-acid sequence, these SPs exhibited inconsistent secretion efficiencies across different cargo proteins. Because the activity of individual SP-cargo pairs is not readily predictable, screening and optimization is often required to achieve target secretion titers. To enhance the utility of our toolbox, we studied the impact of different SP sequence components on protein secretion, then applied this knowledge to develop a standardized toolkit to and enable predictable and tunable protein secretion across diverse SP-cargo pairs. To achieve this, we first identified the lipoprotein export sequence (LES) as the key determinant of efficient secretion of heterologous proteins by lipoprotein SPs. We next performed mutagenesis on the LES region of a representative lipoprotein SP to generate a pool of mutants featuring a standardized SP backbone with diversified LES regions. Screening and characterization of this mutant pool revealed a charge-dependent regulation of both secretion and surface display of heterologous cargo proteins. From these findings, we established a toolkit with improved tunability, enhanced predictability, and surface display capabilities that minimizes the need for iterative screening when developing protein secreting gene circuits for *Bt* and other *Bacteroides* species. By enhancing both the flexibility and control of therapeutic protein output, these results expand the potential of engineered living therapeutic applications, particularly those requiring tunable dosing or surface presentation of proteins.

## Introduction

*Bacteroides* species are among the most abundant and prevalent commensal microbes in the human gastrointestinal (GI) tract, comprising ∼25% of the colonic microbiota *(1)*. The polysaccharide utilization loci (PULs) of *Bacteroides thetaiotaomicron* (*Bt*) account for ∼20% of its genome *(2)*, enabling it to consume a variety of dietary and mucosal polysaccharides *(3)*. This metabolic versatility allows *Bt* to establish long-term colonization in the GI tract, where it is estimated to remain stable for years *(4)*. Furthermore, numerous synthetic biology tools have been developed for the genetic engineering of *Bt (5)*. These features position *Bt* as a promising chassis for developing engineered live bacterial theranostics (eLBTs) for the in situ delivery of biologics and synthetic biology circuits to treat, diagnose, and monitor GI diseases.

Secretion is a primary mechanism for eLBTs to release therapeutic or diagnostic proteins *(6)*. Previously, we identified a set of lipoprotein signal peptides (SPs) that enable high-titer heterologous protein secretion in *Bacteroides* species *(7)*. However, likely due to the diverse folding kinetics, expression levels, and structural stability of different lipoprotein SP-cargo fusion proteins *(8)*, the secretion efficiency of an individual SP cannot be accurately predicted across different cargo protein partners. This limits the ability to precisely control therapeutic dosing, which is critical for balancing efficacy and adverse effects in a clinical setting. There is thus a need to develop secretion tools with enhanced predictability and tunability for *Bacteroides* species.

In this study, we demonstrate that the lipoprotein export sequence (LES) *(9)* is the primary determinant of lipoprotein SP-mediated heterologous protein secretion efficiency in *Bt* and can be engineered for tunability. We show that, when paired with different C-terminal LES, the N-terminal charged (n-) and central hydrophobic (h-) regions of the BT_2479 SP backbone provide a stable scaffold that faithfully reflects the average secretion efficiency mediated by each LES across different lipoprotein SP backbones. Based on this, we constructed a library of BT_2479 SP LES mutants capable of regulating protein secretion from *Bt* at titers spanning three orders of magnitude. Characterization of this library revealed that the highest protein secretion efficiency is achieved with a net charge between +2 to +3 in the LES. We selected 24 LES mutants and one synthetic LES variant to comprise a new secretion toolkit featuring the BT_2479 standardized lipoprotein SP backbone to reduce sequence-dependent impacts on secretion efficiency. We evaluated the ability of these 25 SPs to secrete four distinct heterologous cargo proteins and calculated a “secretion score” for each. Finally, we demonstrated that the BT_2479 SP LES mutants can also regulate protein surface display on *Bt*. This new toolkit, offering enhanced tunability and predictability for both secretion and surface display, significantly improves the utility of *Bt* and expands the potential of eLBT applications.

## Results

### The LES determines the secretion efficiency of lipoprotein SPs

To investigate the factors that govern the heterologous protein secretion efficiency of lipoprotein signal peptides (SPs) in *Bacteroides thetaiotaomicron (Bt)*, we selected four previously reported lipoprotein SPs *(7)* that exhibit diversity in their N-terminal charged (n-), central hydrophobic (h-), and C-terminal LES regions (Fig. 1A). By shuffling the n-, h-, and LES regions of these SPs, we generated a panel of 64 chimeric variants and fused each to the bioluminescent reporter NanoLuc (Nluc) luciferase to quantify secretion efficiency in *Bt* (Fig. 1B). Across all combinations, the LES regions of BT_0294 SP and BT_3740 SP consistently yielded markedly higher and lower Nluc secretion, respectively, regardless of the n- and h-regions they were paired with (Fig. 1B). Notably, the distribution of reporter protein outputs of the 64 variants showed statistically significant differences only when grouped by their LES regions, with the BT_0249 LES consistently demonstrating higher secretion efficiency (Fig. 1C, right), as well as a higher net negative charge (Fig. 1A). In contrast, variants carrying different n-regions exhibited no significant difference in secretion (Fig. 1C, left) despite differences in length and charge (Fig. 1A). Similarly, variants containing the charged BT_3741 h-region (+1) or the longer BT_0294 h-region did not show enhanced secretion relative to variants with other h-regions, all of which are uncharged or 6-7 residues shorter, respectively (Fig. 1C, middle). Together, these results highlight the LES region as the key sequence element controlling the secretion efficiency of lipoprotein SPs and suggest that, beyond the requirement for a charge of at least +1, additional variability in the n- and h-regions does not play a major role in modulating secretion efficiency.

**Figure 1.**
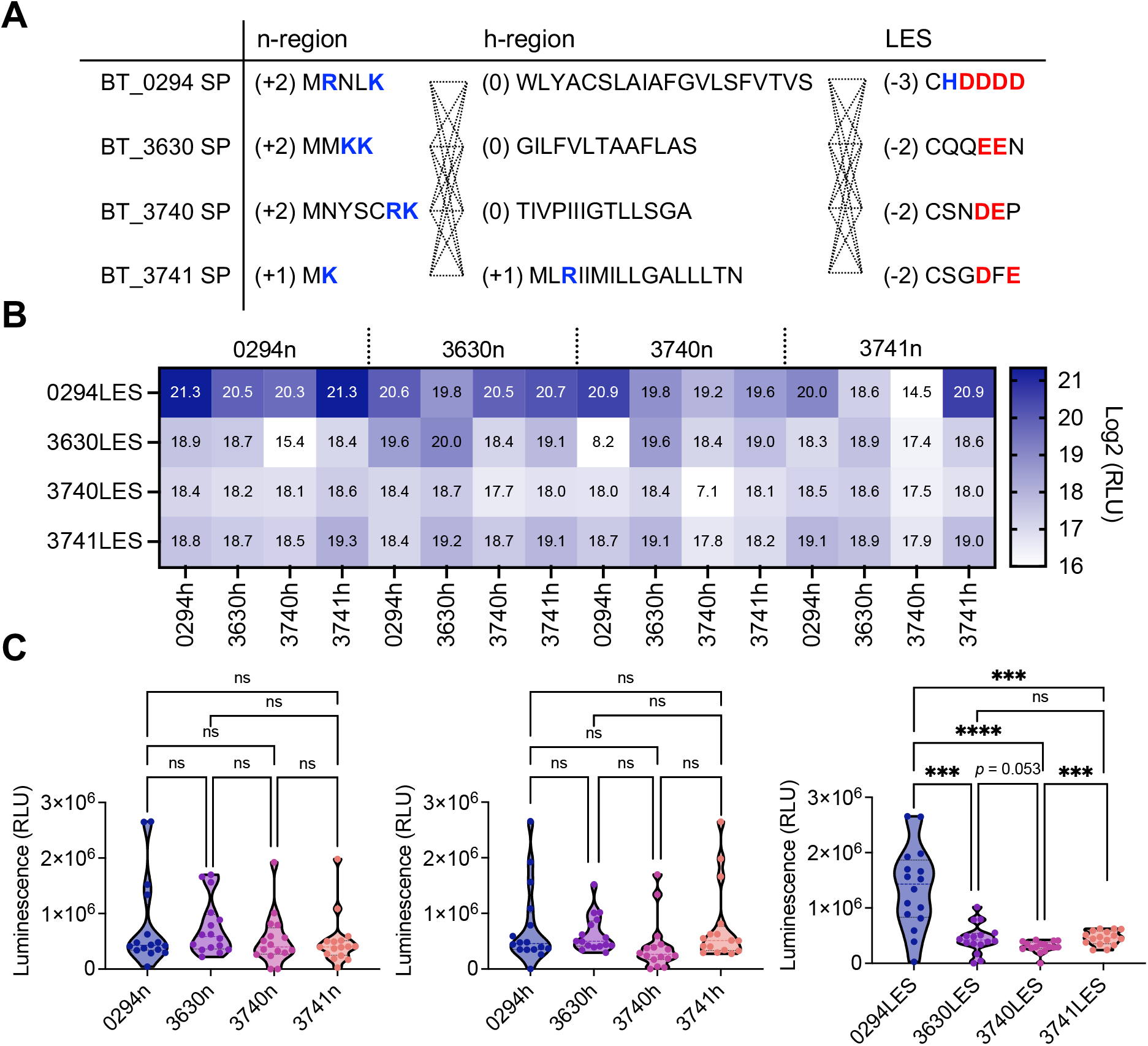
The LES region of lipoprotein SPs drives protein secretion efficiency. (A) Shuffling of n-, h-, and LES regions of four lipoprotein SPs to generate 64 chimeric lipoprotein SP variants. Positively and negatively charged residues are bolded and colored blue or red, respectively, and net charge of each region is shown in parentheses. (B) Secretion of luminescent reporter protein Nluc driven by 64 chimeric lipoprotein SP variants. (C) Protein secretion output grouped based on SP n-, h-, or LES regions. Significance was determined using unpaired two-tailed Welch’s *t* test. ***p < 0.001, ****p < 0.0001; ns, not significant. RLU, relative light unit.

### BT_2479 SP n- and h-regions provide a stable backbone for LES variant testing

While the LES region exerts the strongest influence on secretion efficiency of lipoprotein SPs, the backbone n- and h-regions make minor contributions to this activity (Figs. 1B and 1C), which must be considered for developing a robust and reliable standardized toolkit for tunable secretion. We thus sought to identify a lipoprotein SP backbone which minimally impacts protein secretion efficiency driven by different LES regions.

We selected seven lipoprotein SPs with diverse sequences and shuffled their backbone (n- and h-) and LES regions to generate 49 chimeric variants (Fig. 2A). We measured their secretion efficiency using the Nluc reporter (Fig. 2B) and stratified the luminescence signals by backbone or LES (Fig. 2C). Consistent with our earlier results (Fig. 1C), the protein secretion outputs clustered when grouped by LES rather than by backbone regions. Although this clustering is distinct, we did observe some intra-group variability in secretion efficiency among different LES, potentially arising from minor synergy between the backbone and LES regions.

**Figure 2.**
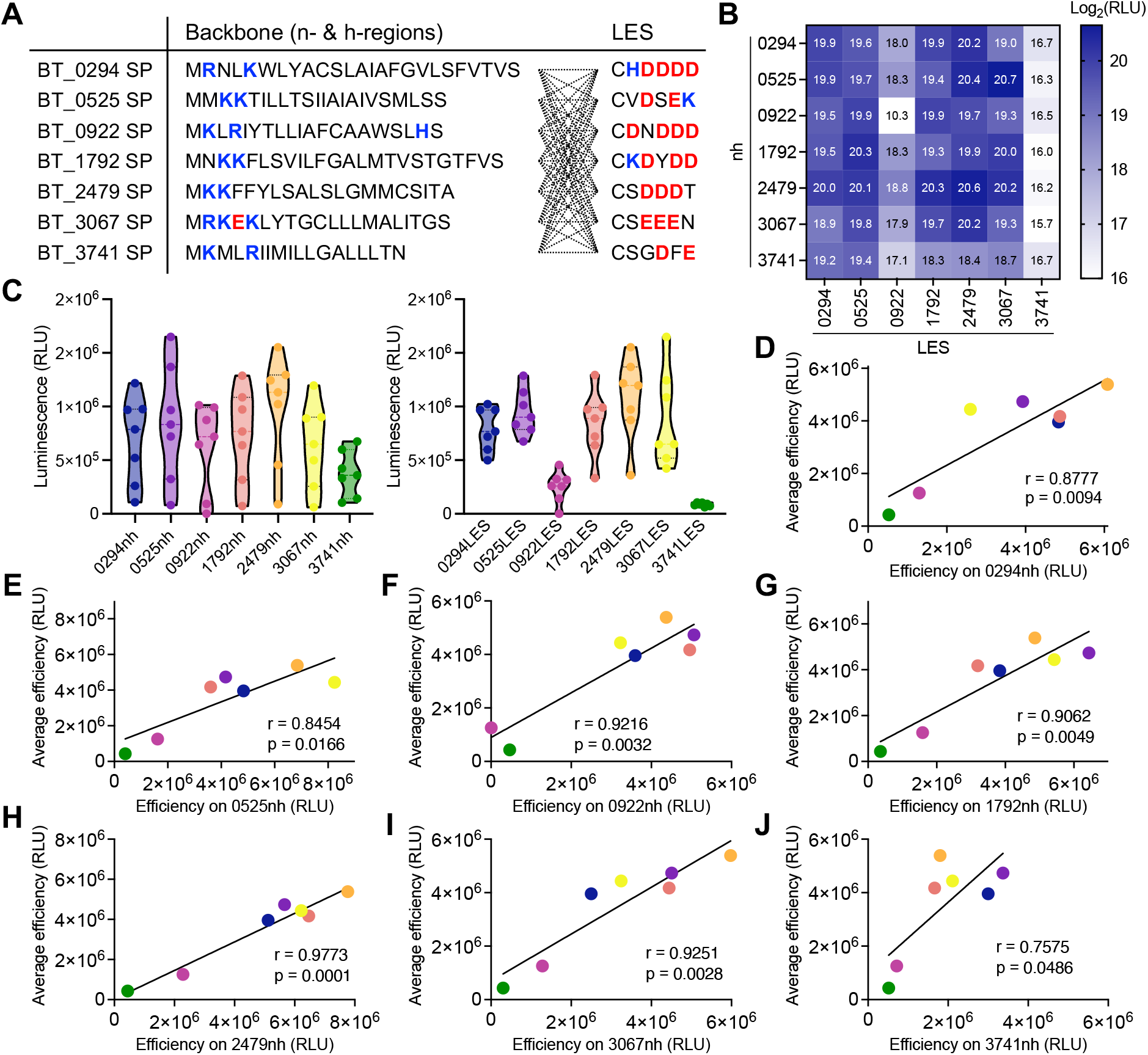
Identification of lipoprotein SP backbone with minimal impact on LES-driven protein secretion efficiency. (A) Shuffling of backbone (n- and h-regions) and LES regions of seven lipoprotein SPs to generate 49 chimeric variants. Positively and negatively charged residues are bolded and colored blue or red, respectively. (B) Protein secretion outputs of Nluc reporter secreted by 49 chimeric lipoprotein SP variants, (C) grouped by backbone (left) or LES regions (right). Data are presented as the mean of triplicate biological samples. (D-J) Correlation between average efficiency of seven LES regions and their individual secretion efficiency when fused with indicated backbone. Data are presented as the mean of triplicate biological samples. The color of each point matches the color of the corresponding LES shown in (C).

To identify the optimal backbone for further development of a standardized SP scaffold, we correlated the secretion efficiencies of the 49 chimeric backbone-LES variants with the average efficiency of the corresponding LES (Figs. 2D-J). The BT_2479 SP backbone exhibited the highest correlation coefficient and statistical significance (Fig. 2H), suggesting that its n- and h-regions exert minimal synergistic effects on LES-dependent secretion efficiency. Accordingly, we selected the BT_2479 backbone as the scaffold for further secretion toolkit development.

### Secretion efficiency of lipoprotein SPs is associated with the net charge of the LES region

Having identified the n- and h-regions of BT_2479 SP as a stable scaffold for measuring LES-mediated protein secretion, we next sought to expand our understanding of the specific determinants of LES activity by characterizing the protein secretion output of a diverse repertoire of LES variants. First, to identify sequence determinants of LES-dependent secretion efficiency, we analyzed the amino acid composition of the LES regions (positions +2 to +6, following the invariant cysteine at position +1) of 109 highly secreted and 37 poorly secreted lipoproteins based on detailed analysis of the presence of native proteins in different fractions of *Bt* culture *(9)* (see Methods). For highly secreted lipoproteins, the SP LES is enriched in aspartate (D), glutamate (E), and asparagine (N), with a strong preference for serine (S) at position +2 (Fig. 3A). Conversely, the LES of SPs from poorly secreted lipoproteins frequently contain lysine (K) and glycine (G) at positions +2, +4, and +5, with no clear sequence conservation at positions +3 and +6 (Fig. 3B).

**Figure 3.**
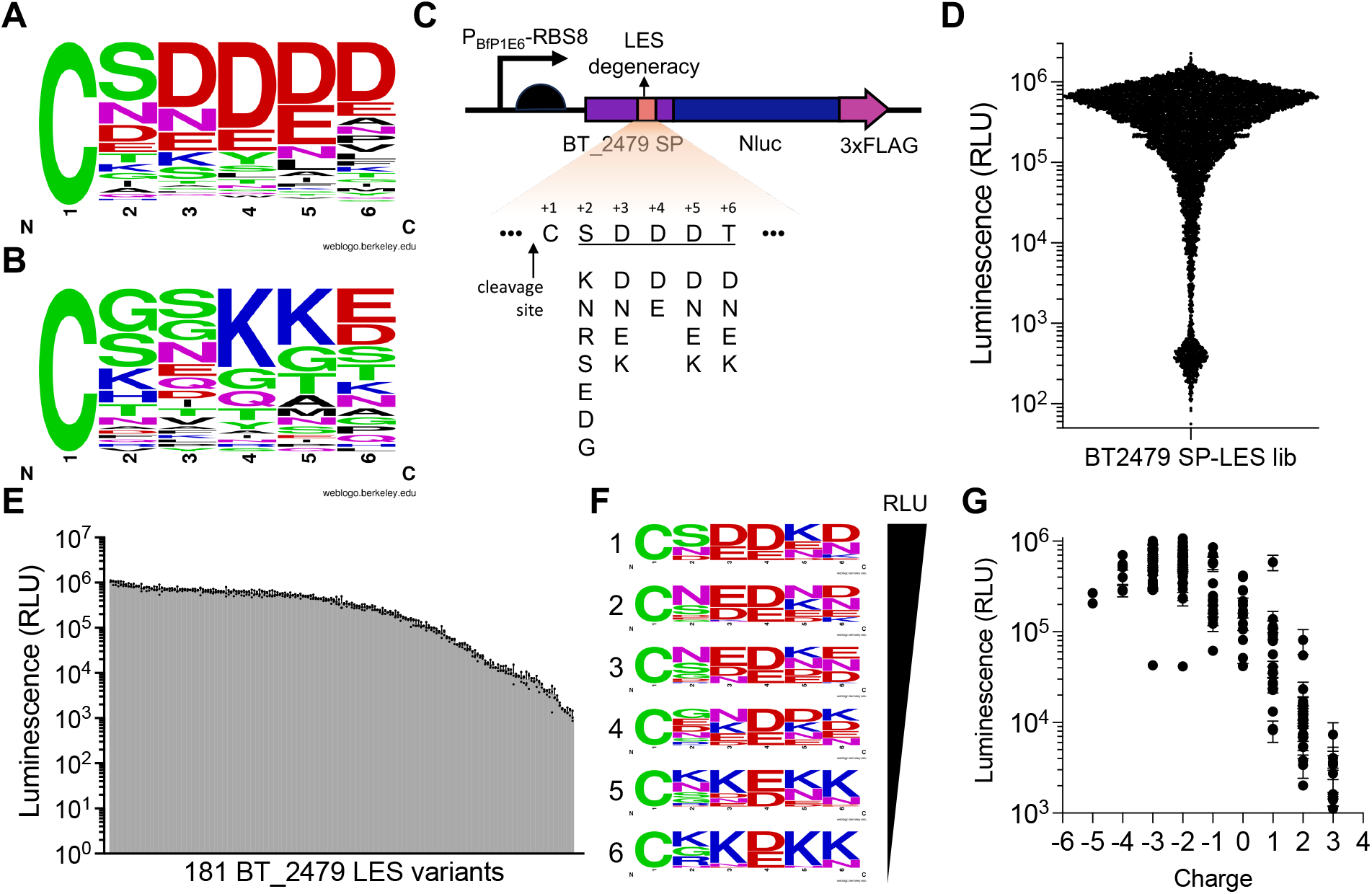
Screening of mutant library reveals optimal charge for LES region. (A and B) Frequency logo plots showing consensus sequences of the six amino acids immediately downstream of the conserved cysteine at the SP cleavage site of (A) 109 highly secreted lipoproteins and (B) 37 poorly secreted lipoproteins native to *Bt*. (C) Design of BT_2479 SP LES mutant library. Expression of Nluc reporter protein was driven by the P_BfP1E6_-RBS8 promoter-RBS pair, which is highly active in *Bt (10)*. A C-terminal 3xFLAG tag was included for immunostaining. (D and E) Protein secretion output, assessed via luminescence measurements of secreted Nluc reporter protein, of (D) all 7,680 mutants from library screening and (E) 181 successfully sequenced mutants. Data in (D) are from single replicates; data in (E) are presented as the mean ± standard deviation of triplicate biological samples. (F) Frequency logo plots of the LES regions of the six sorted groups of the 181 mutants shown in (E), distributed evenly from highest (Group 1) to lowest (Group 6) efficiency of Nluc reporter protein secretion. (G) Nluc reporter protein secretion efficiency plotted as a function of LES net charge of 181 mutants shown in (E). Data are presented as the mean ± standard deviation of triplicate biological samples.

These profiles suggested that the net charge of the LES is a key determinant of secretion efficiency, with the potential to be engineered for tunable secretion. To further refine our understanding of the role of charge in LES-driven protein secretion, we performed site-directed mutagenesis and generated a library of LES mutants of BT_2479 SP. Because the lack of high-throughput screening methods for protein secretion requires manual screening of colonies, we adopted a rational design strategy to maximize targeted diversity while limiting library size. We therefore restricted the library to amino acids that appear to be conserved across LES sequences and are associated with efficient secretion of native Bt lipoproteins (Figs. 3A and 3B). The charged residues aspartate, glutamate, and lysine were incorporated at all positions (Fig. 3C) via the degenerate codon RAM (R: A/G, M: A/C), which also includes the uncharged residue asparagine that is conserved in the LES of highly secreted lipoproteins (Fig. 3A). Moreover, the highly conserved serine and glycine residues (Figs. 3A and 3B) were additionally introduced at position +2 (Fig. 3C) by the degenerate codon RRH (H: A/C/T), which also encodes the conserved but uncharged asparagine (Fig. 3A) and the charged but non-conserved arginine residue. Since position +4 is exclusively aspartate/glutamate or lysine in most native lipoprotein SPs, we reasoned that conservation of these amino acids at this position is critical for secretion and introduction of other amino acids in this location may abolish secretion rather than tuning it. Therefore, to ensure a baseline level of secretion and reduce the number of potentially nonfunctional library members, we restricted position +4 in our library to aspartate/glutamate (degenerate codon = GAM). This design led to a theoretical library size of 1,536 variants.

After conjugating the library into *Bt*, we screened 7,680 colonies (∼five-fold library coverage) using the Nluc reporter to measure secretion output (Fig. 3D). The library exhibited a broad dynamic range of secretion, with signals spanning 10^2^ to 10^6^ relative light units (RLU). We performed direct colony sequencing on 244 colonies from across this range. Notably, nearly all variants with signals <10^3^ RLU contained frameshift mutations in the Nluc coding region and were thus excluded from further analysis, as were samples with sequencing failures, yielding a final count of 181 successfully sequenced BT_2479 SP LES mutant clones. We then individually validated the secretion efficiency of all 181 mutants (Fig. 3E).

To elucidate the sequence-function relationship of the LES domain, we stratified these mutants evenly into six groups based on secretion efficiency and generated logo plots for each (Fig. 3F). This analysis revealed that serine is dominant at position +2 in the top-performing group (Group 1), followed by asparagine (Groups 2–3), and lysine/glycine (Groups 4–6), suggesting that serine at +2 is optimal for high secretion. At position +3, aspartate/glutamate predominated in high-efficiency groups, with a preference for asparagine and then lysine in lower-efficiency groups. A general trend emerged across other positions in which high secretion efficiency correlated with acidic residues (D/E), while low efficiency correlated with basic residues (K). These results underscore the critical role of a negatively charged LES for high-level protein secretion. Notably, the LES regions of the most efficiently secreting mutants (Group 1) were not composed exclusively of negatively charged residues (Fig. 3F), suggesting the existence of an optimal charge window. Indeed, the additive benefit peaked for mutants with a net negative charge of −2 to −3 and diminished once this was increased to −4 and −5 (Fig. 3G).

### BT_2479 SP LES mutants mediate tunable secretion of heterologous proteins via both OMV-dependent and OMV-independent pathways

From the 181 BT_2479 SP LES mutants above (Fig. 3E), we selected 24 mutants with diverse LES net charge (−4 to +3) and Nluc secretion efficiency (10^3^ to 10^6^ RLU) to comprise a new secretion toolkit to enable tunable control of heterologous protein secretion in *Bt*. Additionally, one synthetic LES mutant containing five consecutive lysine residues (KKKKK) with a net charge of +5 was included as a theoretical lower bound of secretion efficiency for this toolkit.

To validate the versatility of the BT_2479 SP LES mutants, we fused these BT_2479 SP LES mutants with three additional cargo proteins: (1) human interleukin 10 (hIL10), a widely used and well-studied anti-inflammatory cytokine *(11, 12)*; (2) a single-domain antibody targeting epidermal growth factor receptor (sdAb-EGFR), found on colorectal and other cancer cells *(13)*; and (3) a single-domain antibody targeting Toxin A (TcdA) of *Clostridium difficile*, a highly destructive gut pathogen *(14)*. We quantified secretion of these three cargo proteins alongside our standard Nluc reporter, driven by the 25 LES mutants noted above, using dot blot analysis. To account for cargo-dependent variation, we calculated a “secretion score” for each mutant based on its aggregated performance across all four proteins (Fig. 4A). Consistent with our earlier findings (Fig. 3G), mutants with a net LES charge of −2 or −3 exhibited the highest secretion scores, which decreased as the net charge increased (Fig. 4A). Although some cargo-dependent variation persisted, the secretion scores were significantly correlated with the individual secretion levels of all four cargoes, with Pearson correlation coefficients (r) ranging from 0.78 to 0.95 (Fig. 4B). These results support the use of this curated set of 25 BT_2479 SP LES mutants as a standardized synthetic biology toolkit for tuning protein secretion efficiency, with the calculated secretion scores serving as reliable predictors of the performance of different LES for novel cargo proteins.

**Figure 4.**
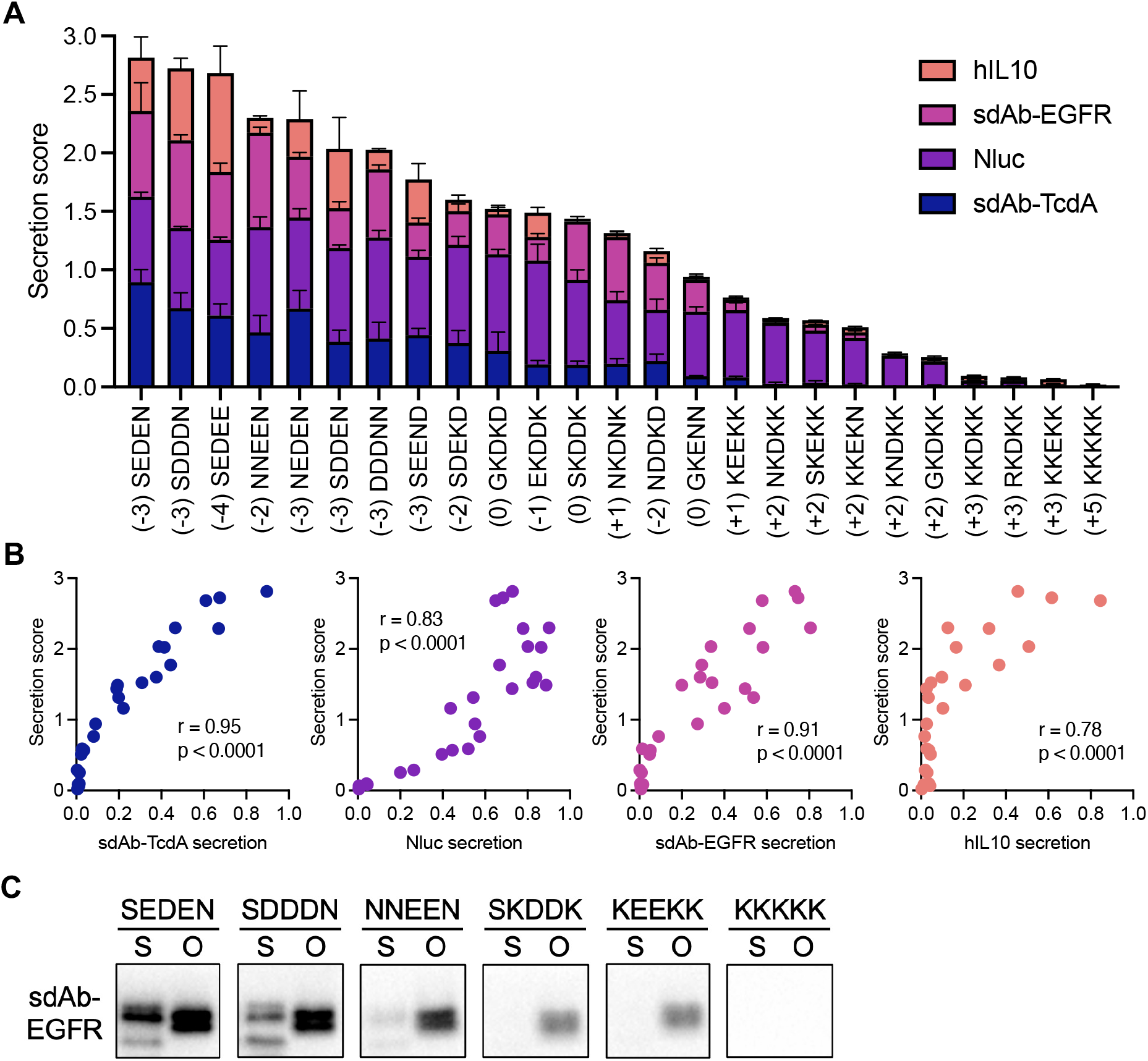
Tunable secretion of heterologous proteins via BT_2479 SP LES mutants. (A) Normalized abundances of secreted cargo proteins and secretion scores of 25 BT_2479 SP LES mutants, with net charge indicated in parentheses. Error bars represent the standard deviation of three biological replicates. (B) Correlation between secretion score of each LES mutant and its normalized secretion level for each of the four cargo proteins. (C) Western blot analysis of protein abundance of sdAb-EGFR secreted by six different LES mutants, measured in the soluble fraction (S) or insoluble OMV fraction (O) of the culture supernatants. OMV fractions were concentrated 20-fold prior to analysis.

Previously, we demonstrated that lipoprotein SPs mediate secretion via both OMV-dependent and OMV-independent pathways *(7)*. To determine whether LES engineering affects one or both pathways, we grew *Bt* cultures expressing the sdAb-EGFR fused downstream of one of six different charge mutants of the BT-2479 SP LES. We then fractionated culture supernatants into soluble (S) and insoluble OMV (O) fractions via ultracentrifugation. Western blot analysis revealed that secretion of both soluble and OMV-associated sdAb-EGFR decreased with increasing LES net charge (Fig. 4C). This suggests that the LES region of the lipoprotein SP regulates secretion through both OMV-dependent and OMV-independent mechanisms.

### BT_2479 SP LES mutants mediate tunable surface display of heterologous proteins

A previous study reported that, in addition to mediating protein secretion via packaging onto OMVs, the LES region may also control lipoprotein export to the outer membrane (OM) *(9)*. We therefore hypothesized that BT_2479 SP LES mutants could also be used for tuning surface display of proteins on *Bt*. To test this, we subjected the *Bt* strains that we previously used to measure protein secretion (Fig. 4A) to flow cytometric analysis to measure surface display of the same cargo proteins. Similar to the secretion score described above, we calculated a “surface display score” for each mutant and observed LES-dependent tuning of protein surface display (Fig. 5A). The display efficiency was strongly associated with the net charge of the LES (Fig. 5A), with rankings closely mirroring those observed for protein secretion (Fig. 4A). Consistent with these results, we observed a high correlation between the secretion scores and surface display scores of BT_2479 SP LES mutants (Fig. 5B).

**Figure 5.**
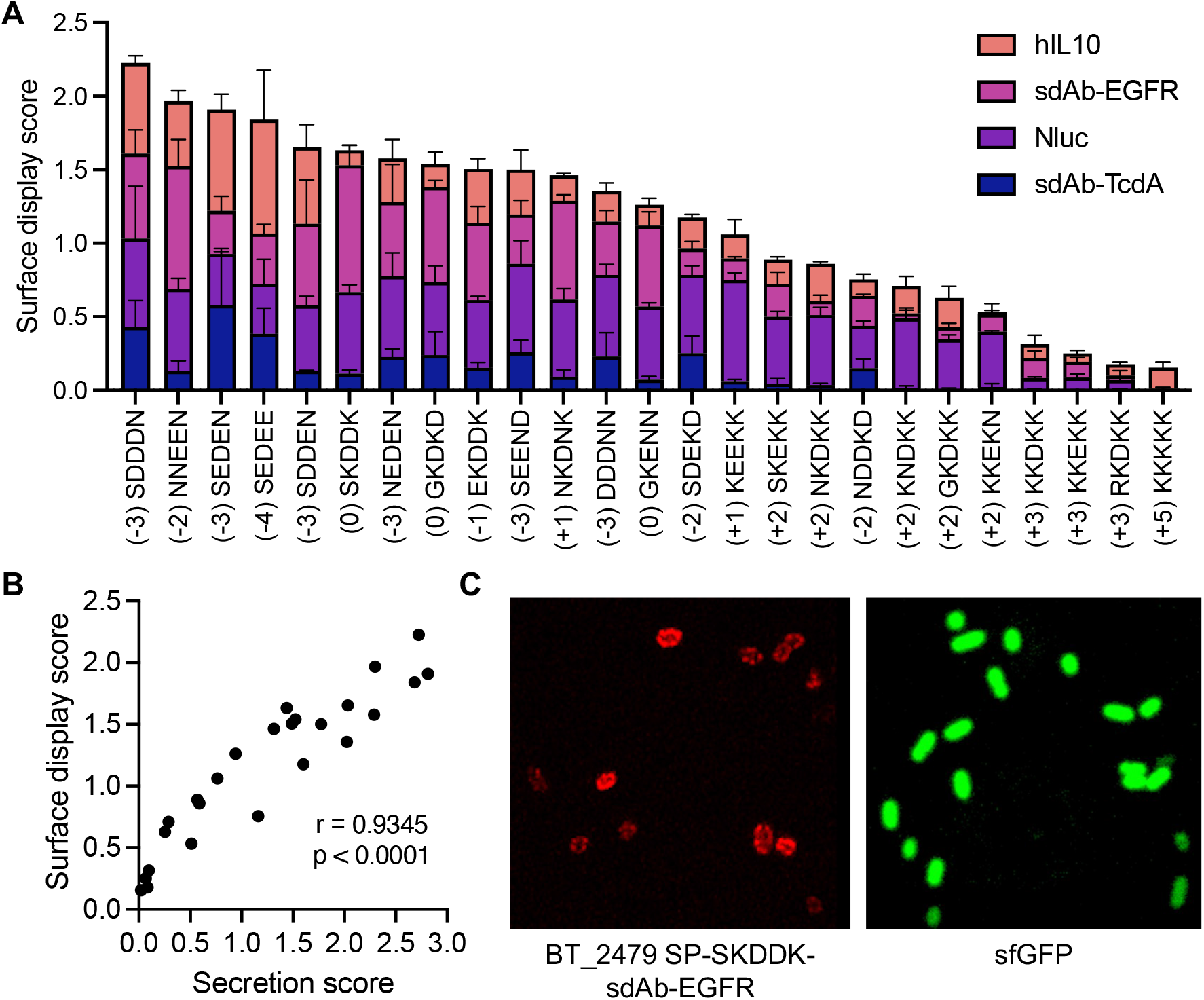
Tunable surface display of heterologous proteins by BT_2479 SP LES mutants. (A) Normalized abundances of surface-displayed cargo proteins and surface display scores of 25 BT_2479 SP LES mutants. Error bars represent the standard deviation of three biological replicates. (B)Correlation between surface display score and secretion score of 25 BT_2479 SP LES mutants. (C) Representative fluorescence microscopy images of (left) *Bt* expressing BT_2479 SP-SKDDK-sdAb-EGFR-3xFLAG stained by anti-FLAG Alexa Fluor Plus 647 or (right) cytosolic sfGFP.

Finally, to confirm surface localization of proteins, we grew *Bt* expressing BT_2479 SP-SKDDK-sdAb-EGFR-3xFLAG, based on its demonstrated high level of surface display (Fig. 5A). Following immunostaining with an anti-FLAG antibody, we analyzed the cells by confocal microscopy and observed fluorescence exclusively at the cell periphery (Fig. 5C, left), demonstrating successful surface display of sdAb-EGFR on *Bt*. By comparison, *Bt* constitutively expressing cytosolic (no SP fusion) superfolder GFP (sfGFP) showed fluorescence evenly distributed throughout the cells (Fig. 5C, right).

## Discussion

*Bacteroides* species are abundant and stable colonizers of the human gut, representing promising vehicles for the long-term, in situ delivery of genetically encoded therapeutics. Secretion is the predominant method for effector protein release in engineered bacteria *(6)*, and our previous work identified various lipoprotein SPs that enable high-efficiency heterologous protein secretion across multiple *Bacteroides* species *(7)*. In that study, however, we observed inconsistent secretion efficiencies, likely due to the variability in length and amino acid composition among these native lipoprotein SPs. In the current work, we more precisely define characteristics of lipoprotein SP-mediated secretion and demonstrate that the lipoprotein export sequence (LES) within the SP determines the efficiency of protein secretion. By performing site-directed mutagenesis on the LES region of the high-efficiency BT_2479 signal peptide, whose backbone (n- and h-regions) exhibits minimal synergy with LES-dependent secretion efficiency, we developed a novel and refined toolkit consisting of standardized BT_2479 LES variants, enabling tunable secretion and surface display of diverse cargo proteins in *Bt*.

It is well established that the charge of the n-region and the length and hydrophobicity of the h-region in Sec signal peptides govern pre-secretion processing, membrane translocation rates, and membrane anchorage in bacteria *(8)*. Our previous findings indicated that a positively charged n-region (net charge ≥ +1) and a sufficiently long h-region (presumably > 12 amino acids) are prerequisites for lipoprotein SP-mediated secretion in *Bt (7)*. Here, we demonstrate that once these basal requirements are met, the charge of the LES becomes the primary determinant of secretion efficiency. Notably, there seems an optimal range of physicochemical properties for each SP region for maximizing protein secretion in different bacteria. For instance, a mutant of the SP for the α-amylase inhibitor tendamistat, with an n-region charge of +2, yielded the highest secretion titers among mutants with n-region charges ranging from +2 to +6 in *Streptomyces lividans (15)*. Likewise, replacing the h-region of the native alkaline phosphatase SP of *E. coli* with a synthetic polyleucine sequence of 10 or 15 residues markedly improved secretion compared with shorter or longer sequences of 5 or 20 residues, respectively *(16)*. Similarly, we found that a net charge within the LES of −2 or −3 maximized secretion of our reporter protein in *Bt*, which is consistent with the results of mutational analyses of the LES of sialidase and mucinase in the *Bacteroidota* member *Capnocytophaga canimorsus (17)*. We speculate that LES charge may influence interactions with unknown transporter proteins during delivery to OMV biogenesis loci on the OM *(18)*, such as the structurally unique β-barrel-assembly machinery (BAM) complex recently revealed and characterized in the phylum *Bacteroidota (19, 20)*. This might parallel the role of the c-region in Sec SPs, which dictates binding specificity for signal peptidase I or II to direct proteins to the periplasm or outer membrane in Gram-negative bacteria *(21)*. However, as the mechanisms underlying lipoprotein transport and OMV biogenesis in *Bacteroides* remain elusive, future studies identifying proteins that interact with lipoprotein SPs will be crucial for elucidating the molecular basis of the optimal physicochemical properties for lipoprotein SP-mediated protein secretion.

Although we standardized the lipoprotein SP backbone, leaving only the LES region variable, the BT_2479 SP LES mutants still exhibited some cargo-dependent variations in secretion and surface display efficiency. While different LES may influence the synthesis and folding rates of specific proteins, other factors such as phenotypic heterogeneity and plasmid copy number variation across selected bacterial colonies may also play a role in these disparities. Indeed, advances in single-cell analysis have highlighted inherent heterogeneity within bacterial populations *(22)*. For example, GFP fluorescence can vary by over two orders of magnitude within an isogenic culture of *Pseudomonas putida (23)*. Furthermore, plasmid copy number can similarly vary among bacterial cells from a single colony of *E. coli*, generating noise in gene expression *(24)*. Consequently, future analyses quantifying single-cell protein expression and plasmid copy numbers may be helpful in deciphering the causes of cargo-dependent variation.

Expanding the concept of secretion to include proteins that remain tethered to the bacterial cell surface further strengthens the therapeutic and diagnostic potential of engineered *Bt*. Indeed, surface display represents a widely adopted strategy for delivering therapeutics via engineered microbes in the GI tract *(6)*, for example: surface display of trefoil factors on *E. coli* has been shown to promote epithelial migration and mucosal repair in colitis *(25)*, and *Lactobacillus plantarum* displaying *Mycobacterium tuberculosis* antigens has been used for oral vaccination *(26)*. More broadly, the availability of surface display tools expands the potential applications of *Bt* across diverse engineering contexts, such as the display of antibody fragments for specific tumor cell targeting *(27-29)*, or the presentation of enzymes for biosensing *(30-33)* and biocatalysis *(34, 35)*. Notably, the simultaneous protein secretion and surface display enabled by the BT_2479 SP LES mutant toolkit described here mirrors well-established protein surface display systems in *E. coli (36)* – such as Lpp-OmpA, autotransporters, and ice nucleation protein – which have also been used to anchor proteins onto secreted outer membrane vesicles (OMVs) *(37, 38)*.

This dual functionality may present both advantages and limitations, depending on the intended application. For applications such as biocatalysis and biodetoxification, having enzymes displayed on the bacterial surface and secreted via OMVs could enhance overall reaction efficiency. However, in applications such as tumor cell targeting, antibody fragments secreted by OMVs may compete with those displayed on the bacterial surface for binding to tumor-associated antigens, diminishing the tumor-targeting efficiency of the bacteria. Notably, two recently described full-length proteins, OmpF (BT_0418) and BT_2844, can display mCherry on the *Bt* surface without concurrent OMV packaging and secretion *(18)*. These proteins may be more suitable for applications requiring cargo retention on the cell surface without release into the surrounding environment. Future investigations into the mechanisms governing the balance between surface display and secretion via OMV packaging will aid in engineering functionally distinct lipoprotein SPs that uncouple these two processes.

In summary, we have developed a standardized lipoprotein SP-based toolkit that provides tunable control over both secretion and surface display of heterologous proteins in *Bt*. By establishing an invariant SP backbone and incorporating secretion scores derived from testing multiple cargo proteins, this toolkit improves predictability and should reduce the screening effort required for new cargo proteins. Furthermore, the tunability of this toolkit expands its versatility for diverse applications, ranging from adjusting therapeutic dose to investigating concentration-dependent, toxin- or enzyme-mediated microbiome-host interactions *(39, 40)*.

## Acknowledgments

This work was funded by NIH NIBIB 5R21EB032548 and S.J.S. is a Biohub Investigator.

## Author contributions

Conceptualization: Y.H.Y. and S.J.S.; Methodology: Y.H.Y.; Formal Analysis: Y.H.Y.; Investigation: Y.H.Y.; Resources: Y.H.Y.; Writing – Original Draft: Y.H.Y. and S.J.S.; Writing – Review & Editing: Y.H.Y. and S.J.S.; Validation: Y.H.Y.; Visualization: Y.H.Y.; Supervision: Y.H.Y.; Project Administration: S.J.S.; Funding Acquisition: S.J.S.

## Materials and Methods

### Bacterial strains and growth conditions

*Bacteroides thetaiotaomicron* VPI-5482 was cultured anaerobically without shaking at 37 °C in Brain Heart Infusion (BHI) medium with the following supplements (BHIS): 1 µg/ml menadione, 0.5 mg/ml cysteine, 0.2 mM histidine, 1.9 mM hematin; or on BHI agar with 10% horse blood (BHIB) as previously described *(7). E. coli* strains were cultured aerobically with shaking in LB medium at 37 °C. Antibiotics were used at the following concentrations: ampicillin 100 µg/mL, kanamycin 50 µg/mL, erythromycin 12.5 µg/mL, chloramphenicol 15 µg/mL, and gentamicin 25 µg/mL for liquid cultures and 200 µg/mL for agar plates.

### Molecular cloning

PCR amplification of DNA was carried out using Q5 high-fidelity DNA polymerase (New England Biolabs). Primers were obtained from Integrated DNA Technologies (IDT). The hIL10 was synthesized (IDT); all other cargo proteins were cloned from previously described plasmids *(7)*. All plasmids were constructed by Gibson Assembly (HiFi DNA Assembly Master Mix, New England Biolabs). Plasmids were transformed into *E. coli* DH5α for storage and maintenance.

### Bacterial conjugation and selection

For each conjugation reaction, 1 mL of overnight culture (∼16-20 h) of *Bt* (plasmid recipient), *E. coli* DH5α (plasmid donor), and *E. coli* RK231 (conjugation helper) were centrifuged at 8,000 x g for 2 min and washed once by PBS. Cell pellets from all three cultures were resuspended together in 30 µL PBS, which was then plated on a BHIB plate and incubated aerobically overnight at 37 °C. The following day, mating lawns were scraped from plates, streaked for single colonies on BHIB plates with 200 µg/mL gentamicin and 12.5 µg/mL erythromycin, and incubated anaerobically for two days.

### Luciferase assay of secreted Nluc

*Bt* transconjugants were inoculated into BHIS medium with 25 µg/mL gentamicin and 12.5 µg/mL erythromycin and incubated anaerobically without shaking at 37 °C overnight. The cultures were centrifuged at 10,000 x g for 3 min to pellet bacteria. Undiluted supernatant was then mixed with 20 µL PBS and 25 µL Nano-Glo^®^ luciferase substrate buffer (Promega). In case of signal saturation, supernatant was diluted 5-10x in PBS prior to proceeding with the PBS/Nano-Glo reaction mixture. Luminescence was measured on a microplate reader using an integration time of 1 s and gain of 100.

### Alignment and visualization of the LES of native secretory lipoproteins of *Bt*

Based on the value of log_2_(OMVs+OMVp/IM+OM) calculated for native secretory proteins *(9)* of *Bt* in our previous study *(7)*, we selected 124 highly secreted lipoproteins (values > 4) and 40 poorly secreted lipoproteins (values < −2). We then analyzed the sequences of these 164 lipoproteins using SignalP 6.0 *(21)* to predict the SP subregions (n-region, h-region, and cleavage site). Several proteins originally annotated as lipoproteins were found to have a Sec/SPI SP or no SP and were thus removed from the candidate set. In addition, we further filtered out the lipoprotein SPs for which the backbone, containing the n- and h-regions, was shorter than 15 amino acids, based on previous findings that such sequences may be incorrectly annotated *(7)*. The remaining 109 highly secreted and 37 poorly secreted lipoproteins were used to create logo plots by WebLogo *(41)*.

### Construction of BT_2479 SP LES mutant library in *E. coli* DH5α

The BT_2479 LES mutant library was constructed using pYHY1-P_BFP1E6_-RBS8-BT_2479 SP-Nluc-3xFLAG *(7)* as the backbone. Mutations were introduced into the LES region using the following primers: BT2479 SP-LES_lib1 R (5’-GATTTTTAGCGTCTATAGTCGTKTYKTYKTCKTYDYYACAAGCCGTAATGGAACACAT CA-3’), BT2479 SP-LES_lib1 insert F (5’-ACGACTATAGACGCTAAAAATCTTGACTATAC-3’), pYHY1 backbone F (5’-GTATGGTGATAGCACACTAGCAC-3’), and pYHY1 backbone R (5’-CTAGTGTGCTATCACCATACTGC-3’). The two DNA fragments amplified by the above primers were assembled by NEBuilder^®^ HiFi DNA Assembly Master Mix, transformed into *E. coli* DH5α. We then scraped ∼16,800 transformant colonies, representing > 10.9-fold coverage of the 1,536-member library, from plates into LB medium. The bacteria were then pelleted by centrifugation at 3200 x g for 10 min, resuspended in 50 mL LB medium with 15% glycerol, and the library was stored in 1.5 mL aliquots at −80 °C.

### Measurement of Nluc secretion efficiency of BT_2479 SP LES mutants in *Bt*

An aliquot of the *E. coli* DH5α BT_2479 SP LES mutant library was thawed and grown overnight in 100 mL LB medium with 100 µg/mL ampicillin. Conjugation mating cultures were mixed, plated, and incubated as described above. The mating lawn was scraped into 30 mL PBS, mixed well by vortexing, and spread on BHIB plates with 200 µg/mL gentamicin and 12.5 µg/mL erythromycin. After two days of anaerobic incubation at 37 °C, we manually picked 7,680 colonies (5-fold coverage of library size) into 96-well plates with 200 µL BHIS containing 25 µg/mL gentamicin and 12.5 µg/mL erythromycin. Following overnight anaerobic incubation at 37 °C without shaking, culture supernatant was separated by centrifugation at 3,200 x g for 10 min, diluted 10-fold in PBS, and 5 µL diluted supernatant was used for Nano-Glo^®^ luciferase assay as described above.

### Dot blot analysis of secreted proteins

*Bt* transconjugants were incubated anaerobically overnight at 37 °C in BHIS medium with 25 µg/mL gentamicin and 12.5 µg/mL erythromycin. The liquid culture was centrifuged at 10,000 x g for 3 min to pellet bacteria and 10 µL supernatant was spotted on a PVDF membrane. The membrane was blocked with 5% milk in PBS-T (0.1% Tween-20 in PBS) for 1 hr at room temperature, then incubated with anti-FLAG M2 monoclonal antibody (Sigma-Aldrich, 1:2000 dilution in 5% PBS-T milk) overnight at 4 °C. The membrane was washed 3 times with PBS-T and incubated with goat anti-mouse IgG secondary antibody conjugated with horse radish peroxidase (HRP) (Jackson Immuno Research, 1:5000 dilution in 5% PBS-T milk) for 1 hr at room temperature. Chemiluminescence was developed using SuperSignal− West Dura Extended Duration Substrate (Thermo Scientific) and detected with a Bio-Rad GelDoc imaging system. The signal intensity was defined as the integrated density of each dot quantified by ImageJ. Signal intensity was further converted into a value between 0 and 1 by min-max normalization, and secretion scores were calculated by summing the normalized signal intensities of four cargo proteins secreted by each BT_2479 SP LES mutant.

### Measurement of surface-displayed proteins

*Bt* strains expressing sdAb-TcdA, sdAb-EGFR, Nluc, and hIL10 fused with each of the 25 BT_2479 SP LES mutants at the N-terminus were inoculated in 96-well plates with 200 µL BHIS containing 25 µg/mL gentamicin and 12.5 µg/mL erythromycin. After overnight anaerobic incubation at 37 °C, 50 µL of liquid culture was transferred into a microcentrifuge tube, pelleted by centrifugation at 8,000 x g for 2 min, and washed once with 400 µL PBS. The cell pellet was resuspended in 50 µL PBS with 1:200 diluted anti-FLAG M2 (Sigma) and 1% bovine serum albumin (BSA) and incubated at 4 °C overnight with rocking. The bacteria were pelleted again, washed once by PBS, and resuspended in 50 µL PBS with 1:100 diluted donkey anti-mouse IgG-Alexa Flour^®^ 647 (Jackson ImmunoResearch) and 1% BSA and incubated at 4 °C overnight with rocking. After pelleting again, the supernatant was removed, bacteria were resuspended in 50 µL PBS, and 10 µL was spotted on slides for fluorescence microscopy (Zeiss LSM 880). Another 30 µL was further diluted into 2 mL PBS for flow cytometry analysis via BD FACSymphony− A1 Cell Analyzer. The fluorescent population (% of 10,000 events) and mean fluorescent intensity were analyzed using BD FACSDiva− software. The abundance of proteins displayed on the surface was calculated as the product of fluorescent population percent and the mean fluorescence intensity. This abundance was then converted into a value between 0 and 1 by min-max normalization. The surface display score was calculated by summing the normalized displayed protein abundance of four cargo proteins for each BT_2479 SP LES mutant.

### Analysis of protein abundance in soluble and insoluble OMV fractions of culture supernatants

*Bt* expressing sdAb-EGFR fused with different BT_2479 SP LES mutants were cultured overnight (20 mL) and centrifuged at 5,000 x g for 10 min to separate cell pellets and supernatants. Supernatants were passed through 0.22 µm syringe filters to remove remaining cells and debris and ultracentrifuged at 150,000 x g at 4 °C for 2 hr to pellet insoluble OMVs. The soluble supernatant fraction (S) was transferred to a new tube. The OMV pellet fraction (O) was washed once with 10 mL PBS, spun down again, and resuspended in 1 mL PBS. The protein concentration of the OMV fraction was measured by DC protein assay (Bio-Rad). The equivalent of 2 µg of protein from the OMV fraction and 25 µL of the soluble fraction were separated by SDS-PAGE, transferred to a PVDF membrane at 100V for 1 hr, and blocked with 5% milk in 0.1% PBS-T (phosphate-buffered saline with 0.1% Tween 20) at room temperature for 1 hr. The membrane was incubated with anti-FLAG M2 monoclonal antibody (1:2000 diluted in 5% PBST-T milk) at 4 °C with rocking overnight. After washing with PBS-T three times, the membrane was incubated with goat anti-mouse IgG secondary antibody conjugated with horse radish peroxidase (HRP) (1:5000 diluted in 5% PBS-T milk) at room temperature for 1 hr. Signal was detected using SuperSignal− West Dura Extended Duration Substrate (Thermo Scientific) on a Bio-Rad GelDoc imaging system.

